# SRSF2 regulation of *MDM2* reveals splicing as a therapeutic vulnerability of the p53 pathway

**DOI:** 10.1101/784256

**Authors:** Daniel F. Comiskey, Matías Montes, Safiya Khurshid, Ravi K. Singh, Dawn S. Chandler

## Abstract

*MDM2* is an oncogene and critical negative regulator of tumor suppressor p53. Genotoxic stress causes alternative splicing of *MDM2* transcripts, which leads to alterations in p53 activity and contributes to tumorigenesis. *MDM2-ALT1* is one of transcripts predominantly produced in response to genotoxic stress and is comprised of terminal coding exons 3 and 12. Previously, we found that SRSF1 induces *MDM2-ALT1* by promoting *MDM2* exon 11 skipping. Here we report that splicing regulator SRSF2 antagonizes the regulation of SRSF1 by facilitating the inclusion of exon 11 through binding at two conserved exonic splicing enhancers. Overexpression of SRSF2 reduced the generation of *MDM2-ALT1* in genotoxic stress condition, whereas knockdown induces the expression of *MDM2-ALT1* in absence of genotoxic stress. Consistently, blocking the exon 11 SRSF2 binding sites using oligonucleotides promotes *MDM2-ALT1*. The regulation of MDM2 splicing by SRSF2 is also conserved in mouse as mutation of one SRSF2 binding site in *Mdm2* exon 11, using CRISPR-Cas9, increases the expression *MDM2-ALT1* homolog *Mdm2-MS2* and proliferation of NIH 3T3 cells. Taken together, these findings underscore the relevance of *MDM2* alternative splicing in cancer and suggest that p53 levels can be modulated by artificially regulating *MDM2* splicing.

## INTRODUCTION

*Murine Double Minute 2 (MDM2)* is a proto-oncogene and critical negative regulator of the p53 tumor suppressor protein. MDM2 is overexpressed in many different types of cancer, including osteosarcoma and other soft tissue sarcomas (1, 2). In response to genotoxic stress *MDM2* undergoes alternative splicing to generate splice variants that are unable to regulate p53 expression (3–5). This in turn results in the stabilization of p53 and subsequent upregulation of pathways involved in apoptosis and cell cycle arrest (6–8). Current targeted therapies to inhibit the interaction between MDM2/p53 in p53 wildtype cancers have had limited success due to toxicity and the fact that they do not also inhibit the regulation of p53 by MDMX. We have previously shown that an isoform of MDM2, MDM2-ALT1 (MDM2-B), can bind and inhibit the proper localization of both full-length MDM2 and MDMX (6). The MDM2-ALT1 isoform is derived from alternative splicing of the full-length *MDM2* pre-mRNA transcript to include only exons 3 and 12. Therefore, understanding the regulation governing this alternative splice variant presents an avenue to stabilize p53 in these cancers for therapeutic benefit.

Alternative splicing is a critical cellular process that generates many transcripts from a single gene. One of the largest families of splicing regulatory proteins is the SR (Serine-Arginine) family of proteins. The role of SR proteins is to enhance or repress the recognition of exons, allowing for increased protein diversity from a single gene (9–12). As a result of this increased proteomic diversity, alternative splicing changes have been associated with multiple cancer cell hallmarks and contribute to tumor progression and therapeutic resistance (13). However, the interplay of protein family members that regulate alternative splicing and contribute to the oncogenic transformation is not well understood.

In order to study the alternative splicing of *MDM2* we have developed a damage-inducible minigene system. The *MDM2* 3-11-12s minigene recapitulates the splicing of the endogenous gene by excluding its intervening exon under genotoxic stress. We have previously identified one of these SR proteins, SRSF1 (also known as ASF/SF2), as a negative regulator of *MDM2* splicing that supports the formation of the *MDM2-ALT1* isoform in response to genotoxic stress (54). SRSF1 binds to an exonic splicing enhancer (ESE) in exon 11 and is necessary to block recognition of exon 11 by the core splicing machinery. In our previous work, we have also identified FUBP1 as another RNA binding protein that regulates *MDM2* splicing. In contrast to SRSF1, FUBP1 binds to an intronic splicing enhancer in intron 11 and facilitates the full-length splicing of *MDM2* (14, 15). However, our *MDM2* 3-11-12s minigene, which lacks the FUBP1 binding site from the full-length *MDM2* 3-11-12, is still efficiently spliced under normal conditions. Therefore, we hypothesize that there are additional sites capable of regulating *MDM2* alternative splicing that can be exploited for therapeutic benefit. Here we report the identification of one such protein, SRSF2 (SC35). We show that mutation of the SRSF2 binding sites or targeting them with splice-switching oligonucleotides (SSOs) increases the expression of the alternatively spliced transcript *MDM2-ALT1*.

## RESULTS

### Mutations in SRSF2 binding sites cause exon exclusion in the *MDM2* 3-11-12s minigene

To better understand the *cis* pre-mRNA sequences and *trans* binding protein factors that facilitate *MDM2* alternative splicing we developed a minigene of *MDM2* that behaves similarly to the endogenous gene under genotoxic stress. We previously reported that the *MDM2* 3-11-12s minigene transcripts undergo exclusion of exon 11 under both UVC and cisplatinum stress (Figure 1A) (15, 16). To identify potential splicing regulator binding sites in the *MDM2* transcript, we scanned the coding sequence of the MDM2 mRNA with ESEfinder (17). Two of the most significant hits were for a pair of SRSF2 binding sites in exon 11 of *MDM2* whose sequences are evolutionarily conserved between mouse and human *MDM2* and flank the previously characterized SRSF1 binding sites (Figure 1B). To determine whether SRSF2 binding sites in exon 11 of *MDM2* affected its alternative splicing, we again turned to ESEfinder to identify changes in specific residues of the SRSF2 binding sites that lowered matrix binding site scores (Figure 1B). We then performed site-directed mutagenesis at one residue of each of the SRSF2 sites as well as created a double mutation by targeting them both simultaneously. We transfected the wild-type and mutant *MDM2* 3-11-12s minigenes into MCF7 cells for 24 hours, then treated under normal (NOR) or UVC conditions, and harvested 24 hours later. RT-PCR analysis revealed that the wild-type *MDM2* 3-11-12s minigene maintains full-length 3.11.12 splicing under normal conditions, and UVC treatment induces the skipped product 3.12. However, mutation of either SRSF2 site (G165T or G213T) in exon 11 resulted in the increased expression of the exon-excluded product in the absence of damage as compared to the wild-type (WT) minigene (Figure 1C, D). Additionally, mutation of both sites together (G165T, G213T) had an additive effect on exon exclusion under normal conditions, indicating that both sites function in the recognition of exon 11 by the spliceosome.

**Figure 1:**
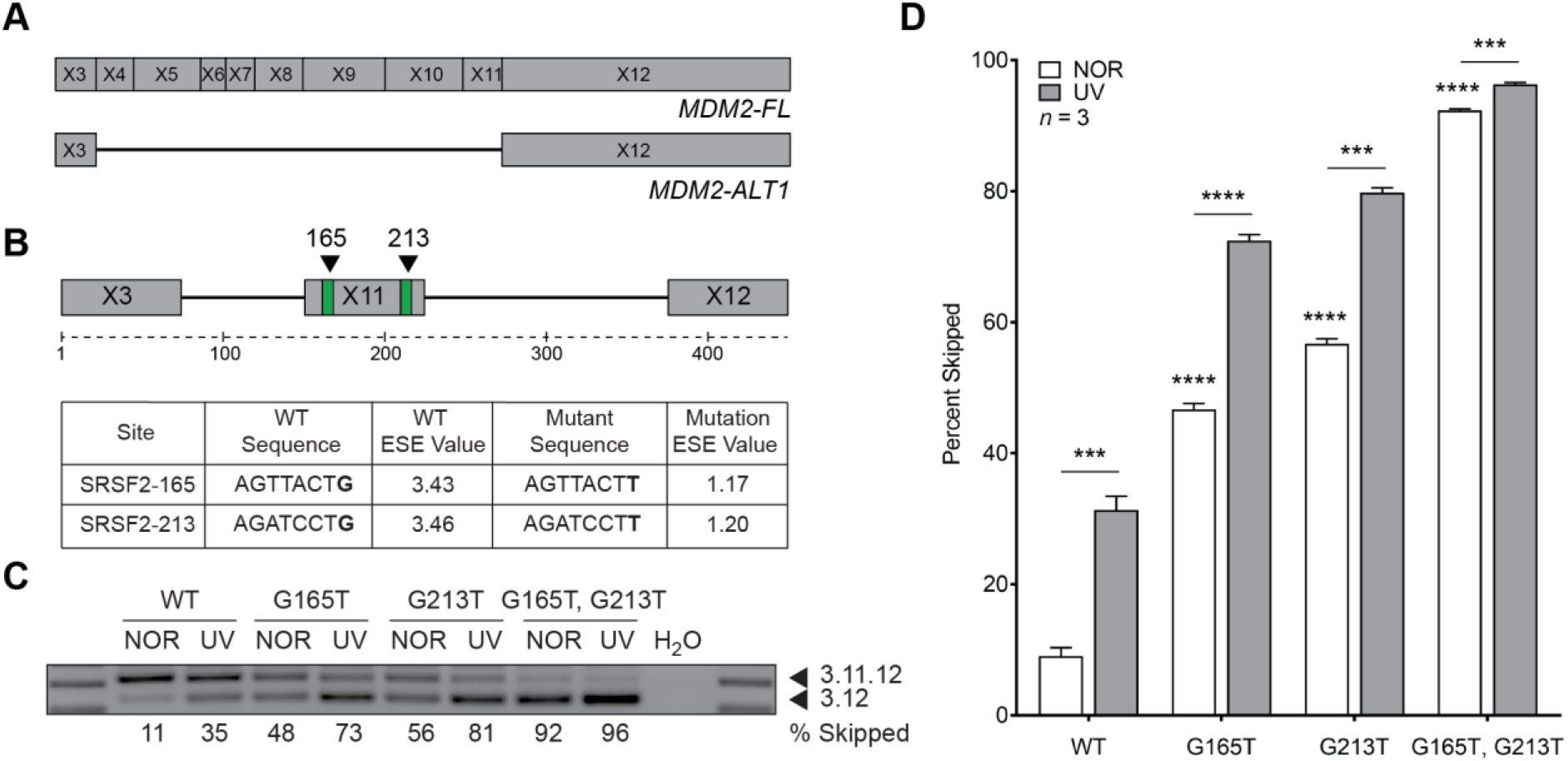
Predicted binding sites for SRSF2 disrupt alternative splicing regulation of the *MDM2* 3-11-12s minigene. **A.** Schematic of full-length *MDM2* and alternative splice variant *MDM2-ALT1.* **B.** Schematic of ESEfinder 3.0-predicted sites for SRSF2 (green) and point mutations made in the *MDM2* 3-11-12*s* minigene (black triangles). Sequences and matrix scores for wild-type (WT) and mutant exonic splicing enhancer (ESE) values were predicted by ESEfinder 3.0. Mutations made in the sequence of the *MDM2* 3-11-12s minigene lowered the predicted matrix score for binding of splicing factor SRSF2. **C.** *MDM2* 3-11-12s minigenes were transfected into MCF7, treated with normal (NOR) or UVC conditions, and harvested 24 hours later. SRSF2 site mutants (G165T *p* = 1.876e-05, G213T *p* = 1.081e-04, G165T, G213T *p* = 3.550e-07) displayed elevated expression of 3.12 under normal conditions compared to the wild-type *MDM2* 3-11-12s minigene. The damage induction of the 3.12 product was maintained in all constructs. **D.** The bar graphs represent the percentage of 3.12 skipped product obtained from three independent experiments under each condition and the error bars represent standard error of the mean (SEM). *** indicates *p* < 0.001, and **** indicates *p* < 0.0001 in all cases as determined by a two-tailed Student’s T test.

### SRSF2 is re-localized in the nucleus and has decreased binding to MDM2 exon 11 in response to UV treatment

Alternative splicing of *MDM2-ALT1* is induced under conditions of genotoxic stress (3). This phenomenon is coincident with an increase in SRSF1 protein expression, a protein we found to cause the induction of MDM2-ALT1 (16). Therefore, we hypothesized that there may be a subsequent decrease in the expression of SRSF2, a protein that supports full-length MDM2 splicing, under the same conditions. We observe that expression of SRSF2 in a nuclear fractionation from HeLa S3 cells increases 12 hours after damage treatment (Figure 2A). Increased SRSF2 levels after stress induction is counterintuitive to our expectations as the expression of *MDM2-ALT1* increases under damage conditions. Therefore, we hypothesized that sequestration of SRSF2 in the nuclear speckles is a potential mechanism of preventing its availability to regulate *MDM2* alternative splicing. To assess SC35 localization is response to damage, we treated MCF7 cells with UV and performed immunofluorescence for SRSF2. Beginning at approximately 4 hours, nuclear speckles became larger and fewer over time (Figure 2C), which correlates with the timing of *MDM2-ALT1* induction (3). We quantified the average size of SRSF2 nuclear speckles before and after 12 hours of damage and found that the average size of nuclear speckle foci was significantly larger after 12 hours of UVC exposure (Figure 2B). These data suggest that SRSF2 relocalization is coincident with *MDM2-ALT1* expression and thus SRSF2 may be playing a direct role in facilitating MDM2 splice site selection in the absence of damage treatment.

**Figure 2:**
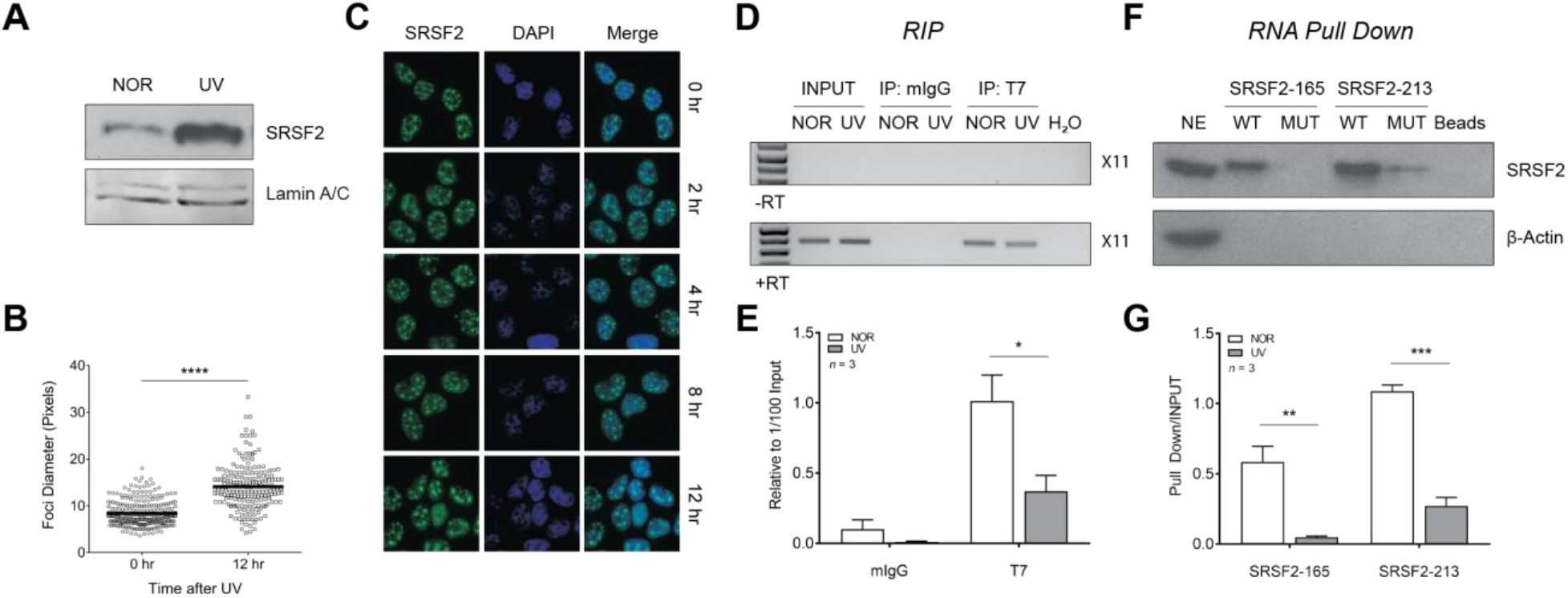
SRSF2 is relocalized and has less affinity for *MDM2* exon 11 upon UV treatment. **A.** SRSF2 protein expression is increased in HeLa S3 in the nucleus after 12 hours of normal or UV treatment. **B.** Quantitation of average nuclear foci diameter at 0 hours and 12 hours after treatment is shown. **C.** Immunofluorescence of SRSF2 in MCF7 shows that SRSF2 (green) was relocalized in the nucleus (DAPI, blue) to fewer and larger nuclear speckles after 12 hours of treatment with 50 J/m^2^ UVC as compared to 0 hours (*p* = 1.71e-47). **D.** RNA immunoprecipitation of T7-SRSF2 and amplification of *MDM2* exon 11. SRSF2 displayed decreased affinity for *MDM2* exon 11 under UV as a compared to normal conditions (*p* = 0.042). Input levels represent RNA levels in 1/100 of immunoprecipitation. Negative isotype (mIgG) and no reverse transcriptase (-RT) controls are also shown. **E.** The bar graphs represent the percentage *MDM2* exon 11 RNA product relative to 1/100 input obtained from three independent experiments under each condition and the error bars represent standard error of the mean (SEM). **F.** RNA oligonucleotides encompassing each SRSF2 binding site, both wild-type (WT) and mutant (MUT) were incubated in nuclear extract (NE) from HeLa S3 cells. Precipitated proteins were washed, then eluted, and run on an SDS-PAGE gel. Mutations in SRSF2 binding sites abrogate binding to each respective site *in vitro* (SRSF2-165 *p* = 0.008, SRSF2-213 *p* = 0.004). Negative ‘beads’ alone control is shown. G. The bar graphs represent the percentage of bound SRSF2 protein relative to nuclear extract input obtained from three independent experiments under each condition and the error bars represent standard error of the mean (SEM). * indicates *p* < 0.05, *** indicates *p* < 0.0001, and **** indicates *p* < 0.0001 in all cases as determined by a two-tailed Student’s T test.

In order to determine whether relocalization of SRSF2 is concurrent with decreased binding to exon 11 of the endogenous *MDM2* pre-mRNA after damage treatment, we performed RNA immunoprecipitation (RIP) with and without damage. SRSF2 bound exon 11 under normal conditions, however this binding was significantly attenuated in conditions of UV stress (Figure 2D and 2E). Furthermore, SRSF2 did not bind to the negative isotype control (mIgG), and we did not observe any amplification of DNA in our control reactions that were not reverse transcribed (-RT). To further demonstrate that ESE mutation leads to abrogated binding of SRSF2 we performed *in vitro* RNA pull down assays using both wild-type and mutant sequences of each binding site. Our results confirm that SRSF2 does in fact bind to both conserved predicted binding sites and mutation of either position significantly abrogates its binding *in vitro* (Figure 2F and 2G) and *in vivo.*

### SRSF2 is a positive regulator of *MDM2* alternative splicing

In order to assess the role of SRSF2 as a positive regulator or *MDM2* alternative splicing we performed overexpression and knockdown experiments. We began by co-transfecting MCF7 cells with the *MDM2* 3-11-12s wild-type minigene along with either LacZ as a negative control, or T7-SRSF2, followed by mock or UV treatment (50J/m^2^ UVC). We observed that transfection of SRSF2 abolished damage-responsive alternative splicing of minigene transcripts under UVC conditions as compared to the negative control (Figure 3A, B). Conversely, we treated MCF7 cells with siRNA against SRSF2 and confirmed its efficient knockdown (Figure 3C). We then performed a nested RT-PCR to identify transcripts from the endogenous *MDM2* gene. We found that *MDM2-ALT1* (3.12) was significantly induced in the absence of any genotoxic stress as compared to no treatment (NT) and control siRNA (CTRL) (Figure 3D). These data indicate that SRSF2 expression is required to facilitate inclusion of all *MDM2* internal exons (exons 4 through 11), not only the penultimate exon 11 as studied in our minigene system.

**Figure 3:**
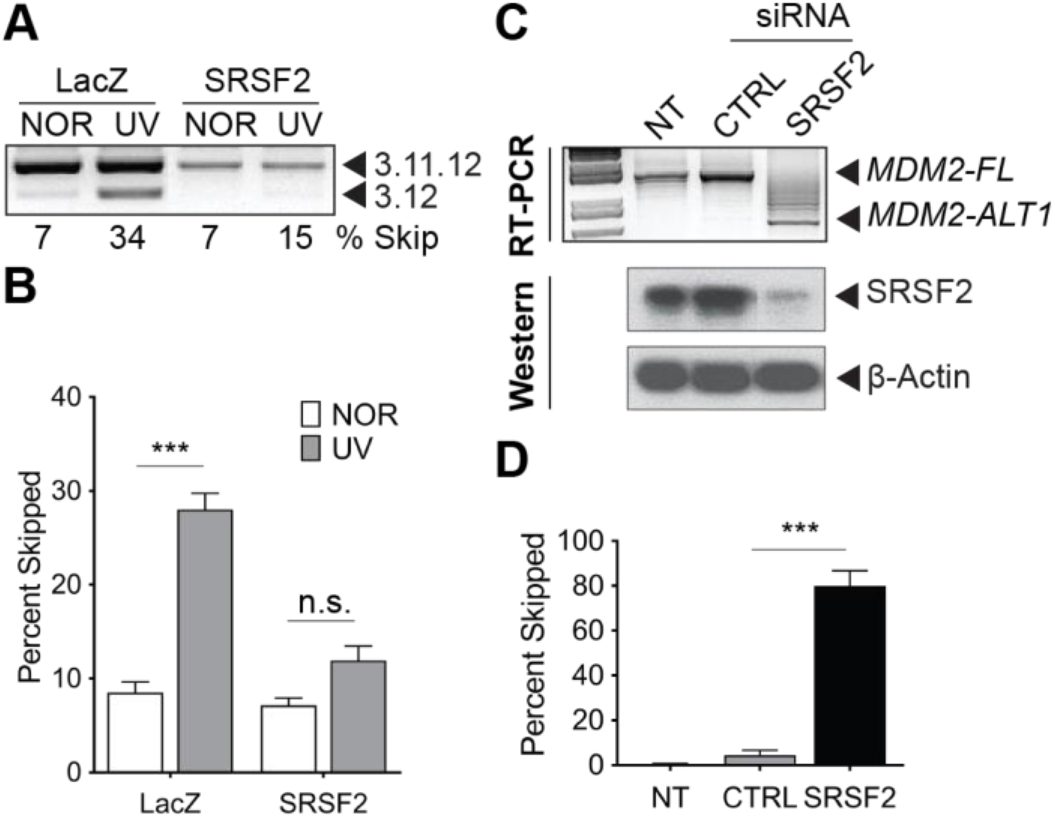
SRSF2 is a positive regulator of *MDM2* splicing. **A.** RT-PCR analysis of MCF7 cells cotransfected with the wild-type *MDM2* 3-11-12s minigene and a negative control, LacZ, or SRSF2 expression construct. Transfection of SRSF2 ablated damage-induced alternative splicing of the wild-type *MDM2* 3-11-12s minigene. (*p* = 1.312e-05 LacZ, *p* = 0.050 SRSF2). **B.** The bar graphs represent the percentage of 3.12 skipped product obtained from five (LacZ) or three (SRSF2) independent experiments under each condition and the error bars represent standard error of the mean (SEM). **C.** MCF7 cells were either non-transfected (NT) or transfected with either a control (CTRL) or gene-specific siRNA (SRSF2) for 72 hours. A representative nested RT-PCR and Western blot are shown. Beta-actin was used as a loading control. Knockdown of SRSF2 significantly induced the alternative splicing of endogenous *MDM2-ALT1* in the absence of damage (*p* = 4.668e-04). **D.** The bar graphs represent the percentage of *MDM2-ALT1* skipped product relative to *MDM2-FL* obtained from three independent experiments under each condition and the error bars represent standard error of the mean (SEM). The error bars represent standard error of the mean (SEM). *** indicates *p* < 0.001 in all cases as determined by a two-tailed Student’s T test.

### Splice-switching oligonucleotides (SSOs) against SRSF2 sites in exon 11 induce expression of *MDM2-ALT1*

We hypothesized that SSOs targeting SRSF2 binding sites would occlude SRSF2 binding and could therefore be used to induce skipping the internal exons of *MDM2.* To test this hypothesis, we designed SSOs against each of our identified binding sites (Figure 4A). Briefly, we co-transfected the wild-type *MDM2* 3-11-12s minigene along with SSOs against SRSF2 sites for 24 hours in MCF7 cells. RT-PCR of exogenous *MDM2* revealed that SSOs against either SRSF2-165 (SSO1) or SRSF2-213 (SSO2, SSO3) site were effective at inducing expression of the exon-excluded product 3.12 under normal conditions compared to non-specific SSO (NS-SSO; Figure 4B). Whereas SSO1 against the first SRSF2 site induced 3.12 modestly at a range of concentrations, SSOs 2 and 3 against the second SRSF2 site were far more potent in inducing exon exclusion at all concentrations (Figure 4C). These data are consistent with mutation of the SRSF2 binding sites in the *MDM2* minigene described above.

**Figure 4:**
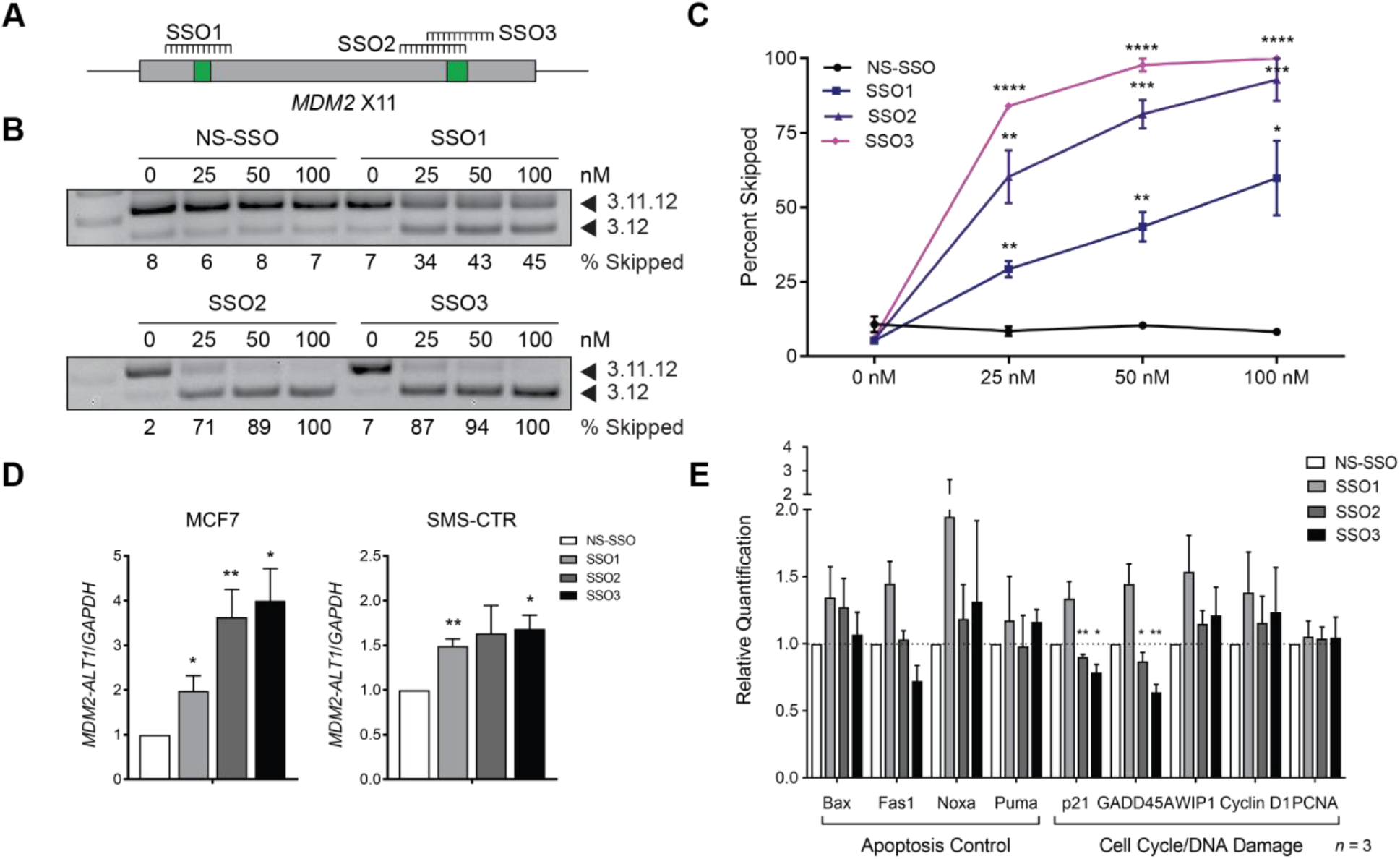
Splice-switching oligonucleotides (SSOs) targeting SRSF2 sites in *MDM2* exon 11 induces expression of *MDM2-ALT1.* **A.** Schematic of binding site for SSOs targeting SRSF2 sites in *MDM2* exon 11. **B.** The wild-type *MDM2* 3-11-12s minigene was cotransfected with either 0 nM, 25 nM, 50nM, 100 nM of each indicated SSO into MCF7 cells and harvested 24 hours later. MCF7 cotransfected with SSOs against SSO1, SSO2, and SSO3 showed increased exon skipping compared to NS-SSO at all concentrations. **C.** The bar graphs represent the percentage of 3.12 skipped product obtained from three independent experiments under each condition and the error bars represent standard error of the mean (SEM). **D.** Cells were transfected with 100 (MCF7) or 250 nM (SMS-CTR) non-specific (NS-SSO) or SRSF2 site-specific SSOs (SSO1, SSO2, SSO3) for 24 hours and subjected to qRT-PCR for *MDM2-ALT1* and normalized to *GAPDH.* SSO1, SSO2, and SSO3 induced expression of *MDM2-ALT1* as compared to NS-SSO in MCF7 cells (n = 4, *p* = 0.028 SSO1, *p* = 0.006 SSO2, *p* = 0.014 SSO3) and SMS-CTR cells (n = 3, *p* = 0.017 SSO1, *p* = 0.011 SSO3). E. MCF7 cells were transfected with either 100 nM NS-SSO, SSO1, SSO2, or SSO3 for 24 hours and extracted cDNA was subjected to a qRT-PCR for p53 target genes and normalized to *GAPDH.* Transfection of SSO2 and SSO3 significantly reduced the expression of *GADD45A* and *CDKN1A* compared to NS-SSO. The error bars represent standard error of the mean (SEM). * indicates *p* < 0.05, and ** indicates *p* < 0.01 in all cases as determined by a two-tailed Student’s T test.

Decreasing the levels of SRSF2 caused skipping of multiple internal exons of the endogenous *MDM2* transcript, generating *MDM2-ALT1*. We therefore hypothesized that SSOs targeting the SRSF2 binding sites in exon 11 could likewise induce MDM2-ALT1 expression. We transfected either NS-SSO, SSO1, SSO2, or SSO3 into MCF7 or SMS-CTR cells. We then performed a qRT-PCR assay specific for *MDM2-ALT1*, which targets the splice junction between exons 3 and 12. Consistent with our results with *MDM2* 3-11-12s minigene we were able to observe a significant increase in expression of *MDM2-ALT1* (Figure 4D).

We assessed the functional impact of SSO treatment on the p53 pathway by examining transcriptional target levels, as well as cell cycle changes. We found that SSOs targeting the second SRSF2 site (SSO2, SSO3) significantly reduced the expression of *GADD45A* and *CDKN1A* in MCF7 cells (Figure 5E). We also performed cell cycle analysis but did not find any changes in the any phase of the cell cycle between NS-SSO, and SSO1, SSO2, or SSO3-transfected MCF7 cells (data not shown). MCF7 cells are *ARF* null and therefore lack an intact p53 pathway (18), which may explain the lack of induction of p53 in response to MDM2-ALT1 expression. Additionally, given the transient nature of our transfection and the modest level of *MDM2-ALT1* induction, it is not surprising that there were no overt phenotypic changes in response to SSO treatment.

**Figure 5:**
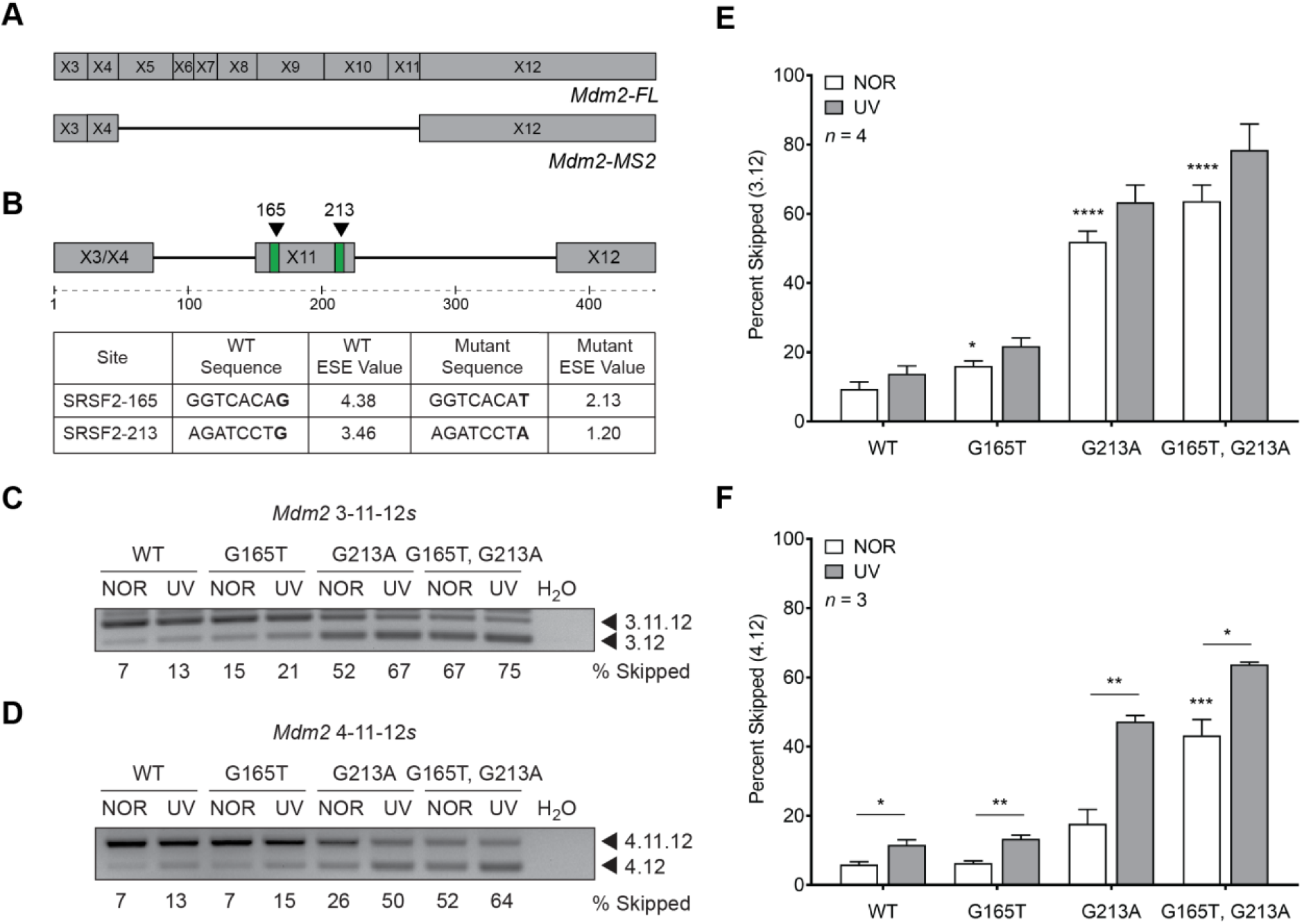
Conserved SRSF2 binding sites disrupt alternative splicing of mouse *Mdm2.* **A.** Schematic of full-length *Mdm2* and alternative splice variant *Mdm2-MS2.* **B.** Schematic of *Mdm2* exon 11 with ESEfinder 3.0 predicted sites for SRSF2 (green) and point mutations made in the *Mdm2* 4-11-12s minigene (black triangles). Sequences of SRSF2 binding sites and matrix scores for wild-type (WT) and mutant exonic splicing enhancer (ESE) values that were predicted by ESEfinder 3.0. Mutations made in the sequence of the *Mdm2* 4-11-12s minigene lowered the predicted matrix score for binding of splicing factor SRSF2. **C.** *Mdm2* 4-11-12s minigenes were transfected into mouse myoblast C2C12 cells in order to better assess their native context, treated under normal (NOR) or UVC conditions, and harvested 24 hours later. SRSF2 mutants (G165T, G213A, *p* = 1.356e-03) displayed increased expression of 4.12 under normal conditions compared to the wild-type *Mdm2* 4-11-12s minigene. The damage induction of the 3.12 product was maintained in all constructs. **D.** *Mdm2* 3-11-12*s* minigenes were transfected into MCF7 cells for 24 hours, then treated under normal (NOR) or UVC conditions, and harvested 24 hours later. SRSF2 mutants (G165T *p* = 0.038, G213A *p* = 2.451e-05, G165T, G213A *p* = 3.910e-05) displayed increased expression of 3.12 under normal conditions compared to the wild-type *Mdm2* 3-11-12s minigene. **E.** The bar graphs represent the percentage of 3.12 skipped product obtained from three independent experiments under each condition and the error bars represent standard error of the mean (SEM) **F.** The bar graphs represent the percentage of 3.12 skipped product obtained from four independent experiments under each condition and the error bars represent standard error of the mean (SEM). * indicates *p* < 0.05, ** indicates *p* < 0.01, *** indicates *p* < 0.001, and **** indicates *p* < 0.0001 in all cases as determined by a two-tailed Student’s T test.

### SRSF2 regulation of *MDM2* splicing is conserved from mouse to human

Like the human *MDM2* gene, the mouse *Mdm2*, undergoes alternative splicing under conditions of genotoxic stress. The resultant predominant splice variant, *Mdm2-MS2*, however, is different than the human MDM2-ALT1 and comprises exons 3, 4, and 12 (Figure 5A) (3). To determine the conserved regulation of *MDM2* splicing by SRSF2 regulation, we generated a mouse minigene that contains the *Mdm2* exon 3 as the first exon joined to exon 11 and exon 12, as in the human construct. We next examined the sequence of mouse exon 11 to identify mutations that lower the ESE matrix scores for predicted SRSF2 binding sites in the mouse gene. Mutations similar to those we made in the human minigene lowered the predicted binding scores of the ESE in the mouse Mdm2 exon 11 (Figure 5B). We then induced the G165T or G213A mutations, or both, into the mouse minigene and tested their ability to support Mdm2 splicing regulation. Mutations of both the G165T and G213A residues resulted in increased expression of exon 11-excluded product in the absence of genotoxic stress (Figure 5C, E). As with the *MDM2* 3-11-12s G213T mutation, the *Mdm2* 3-11-12s G213A mutation was more potent than the G165T mutation. It should be noted, however, that the mouse minigene does not accumulate increased exon 11 skipping as a result of UV treatment.

**Figure 6:**
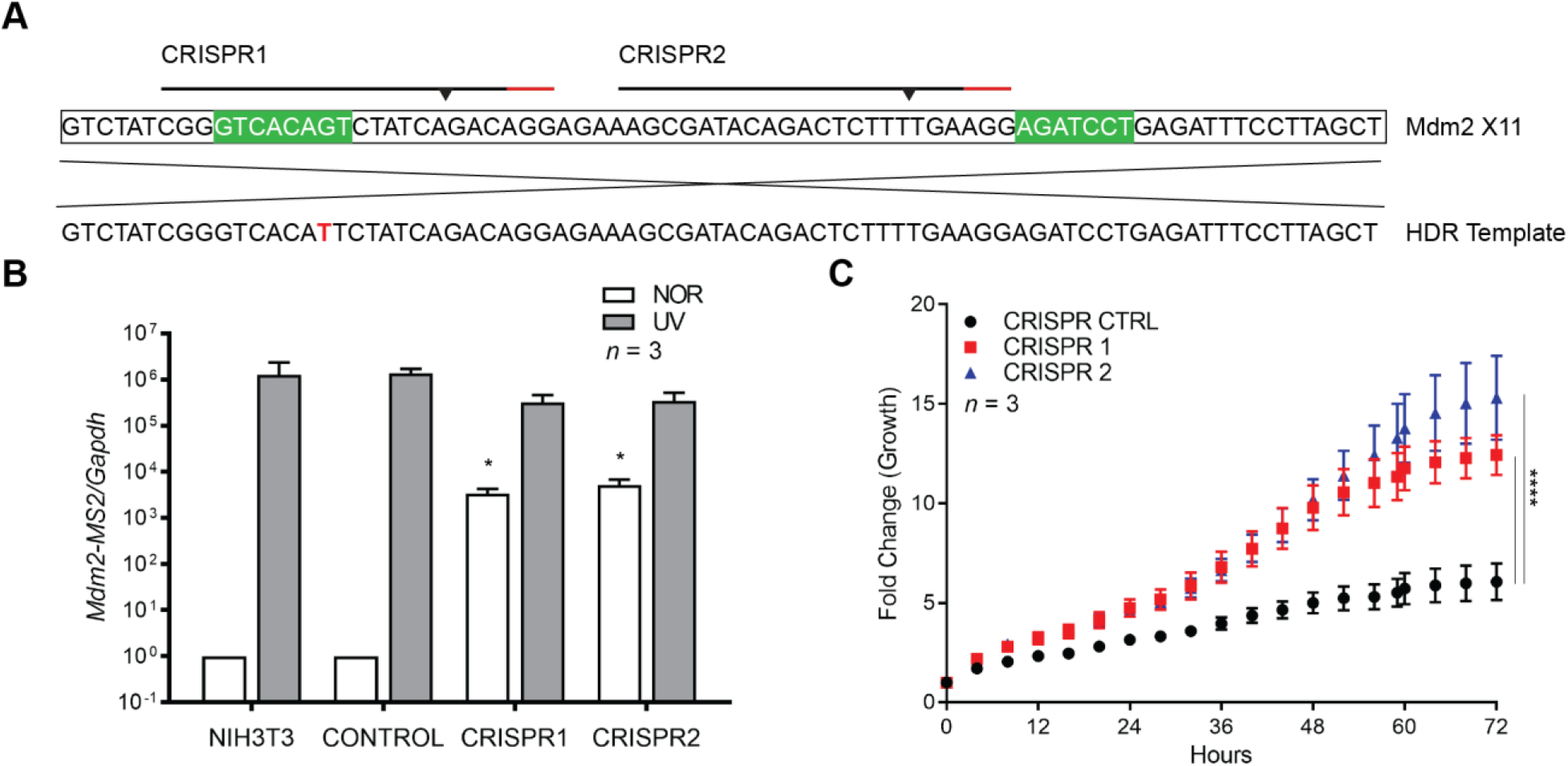
Endogenous mutation of SRSF2 binding site in *Mdm2* increases alternative splicing and cell proliferation. **A.** CRISPR sites in mouse *Mdm2* exon 11 (black lines), with protospacer adjacent motifs (red lines), predicted double-strand break sites (triangles), and SRSF2 binding sites (green). The homology-directed repair (HDR) template is displayed below with SRSF2 splicing mutation (red). **B.** NIH 3T3 cells wild-type or mutant for SRSF2-G165T were treated under normal (NOR) or UVC conditions, and harvested 24 hours later. Cells were harvested and a qRT-PCR was performed for *Mdm2-MS2.* Engineered SRSF2 splice mutant cell lines (CRISPR1 *p* = 0.021, CRISPR2 *p* = 0.037) showed increased levels of *Mdm2-MS2* in the absence of damage compared to NIH 3T3 wild-type cells. **C.** NIH 3T3 cells wild-type (CONTROL) or SRSF2-G165T mutants (CRISPR1, CRISPR2) were seeded in a 6-well plate. Cell density was measured in a cell confluence assay using an IncuCyte live-cell analysis system every four hours for three days. SRSF2 splice mutants (CRISPR1 *p* and CRISPR2) showed significantly increased proliferation at 72 hours compared to wild-type control (CONTROL). The graphs represent fold change in growth obtained from twelve independent experiments under each condition and the error bars represent standard error of the mean (SEM). * indicates *p* < 0.05, and **** indicates *p* < 0.0001 in all cases as determined by a two-tailed Student’s T test.

Because the mouse splice variant, *Mdm2-MS2*, is comprised of exon 4 spliced directly to exon 12, we wondered if the sequences in Mdm2 exon 4 may be important to achieve regulation in response to damage treatment in the mouse gene. We engineered a second mouse minigene containing exons 4, 11 and 12 and induced these same mutations in an *Mdm2* 3-11-12s minigene to assess in cell culture with and without UV damage. We observed that both G165T and G213A mutations together resulted in an increase of exon skipping under normal conditions (Figure 5D, F), and were induced by UV treatment. The effects of the G165T and G213A mutations were additive in both the 3-11-12s and 4-11-12s minigene, suggesting that these SRSF2 sites regulate *Mdm2* splicing independently.

### *CRISPR-Cas9-engineered* mutant cell lines demonstrate endogenous regulation of *Mdm2* by SRSF2 sites in exon 11

To precisely pinpoint the regulation of SRSF2 on *Mdm2* endogenous transcripts we used CRISPR-Cas9 to induce SRSF2 site mutations. We designed Cas9 guide RNAs (gRNAs) to exon 11 of *Mdm2* as well as single-stranded oligonucleotide donor (ssODN) repair templates that included the nucleotide mutation at our sites of interest (Figure 6A). Our attempts to recover cells with both mutations were unsuccessful, so we designed separate 243 base pair ssODNs that contained a single point mutation, targeting each site independently. Eventually, we recovered NIH 3T3 cell lines with the single G165T mutation but were unable to generate cell lines with the G213A mutation. We subjected both control NIH 3T3 cell lines transfected with a control CRISPR plasmid and cells bearing the G165T mutation to a qPCR to specifically detect expression of *Mdm2-MS2.* We report that cells with the G165T mutation demonstrated a significantly higher amount of *Mdm2-MS2* under normal conditions as compared to control cells (Figure 6B).

To test the effect of *Mdm2-MS2* expression on cell proliferation, we seeded wild-type control CRISPR cells, and both SRSF2 G165T-engineered cell lines (CRISPR1, CRISPR2) and monitored them continuously for growth. At the end of 72 hours both of our G165T cell lines showed significantly higher fold change in cell confluence (CRISPR1 and CRISPR2) and hence increased proliferation in comparison to the wild-type CRISPR control (CRISPR CTRL, Figure 6C). It is important to note that NIH 3T3, like MCF7 cells, though wild-type for p53, are mutant for *ARF.* In the absence of an intact p53 pathway, MDM2 splice variants will have p53 independent functions that support increased cell proliferation and transformation. Indeed, it has been demonstrated by others that *MDM2* splice variants are capable of transforming NIH 3T3 cells (19).

### SRSF1 and SRSF2 are antagonistic for the control of *MDM2* splicing

Given that the identified SRSF2 sites flank the previously identified SRSF1 binding site, we wanted to test the dependency of these sites in counterbalancing the activity of the other. We hypothesized that the positive action of the SRSF2 binding is required to overcome the negative splicing function of the SRSF1 binding. We induced the previously published SRSF1 mutation together with one or both of the SRSF2 mutations and tested them in our damage-inducible cell culture system (Figure 7A). We observed that both the SRSF2-G165T and the SRSF2-G213T mutation were sufficient to overcome the mutation of the SRSF1 sites, and restore the damage induction of exon 11 skipping (where none was seen in SRSF1 mutant minigene). Lastly, when both SRSF2 sites were mutated in the context of the SRSF1 mutant we again observed a significant increase in the level of exon skipping under normal conditions (Figure 7B and 7C). These data strongly suggest that the regulation of *MDM2* alternative splicing by SRSF2 is necessary to counteract the negative effects of SRSF1. These data further suggest that the density of splicing elements and functional redundancy of regulators in this region plays an important role in the alternative splicing of *MDM2* since splicing control is maintained in the absence of both the SRSF1 and SRSF2 elements. Furthermore, identification of additional elements will allow improvement of splice altering therapies that could be used to modulate the p53 pathway in cancer.

**Figure 7:**
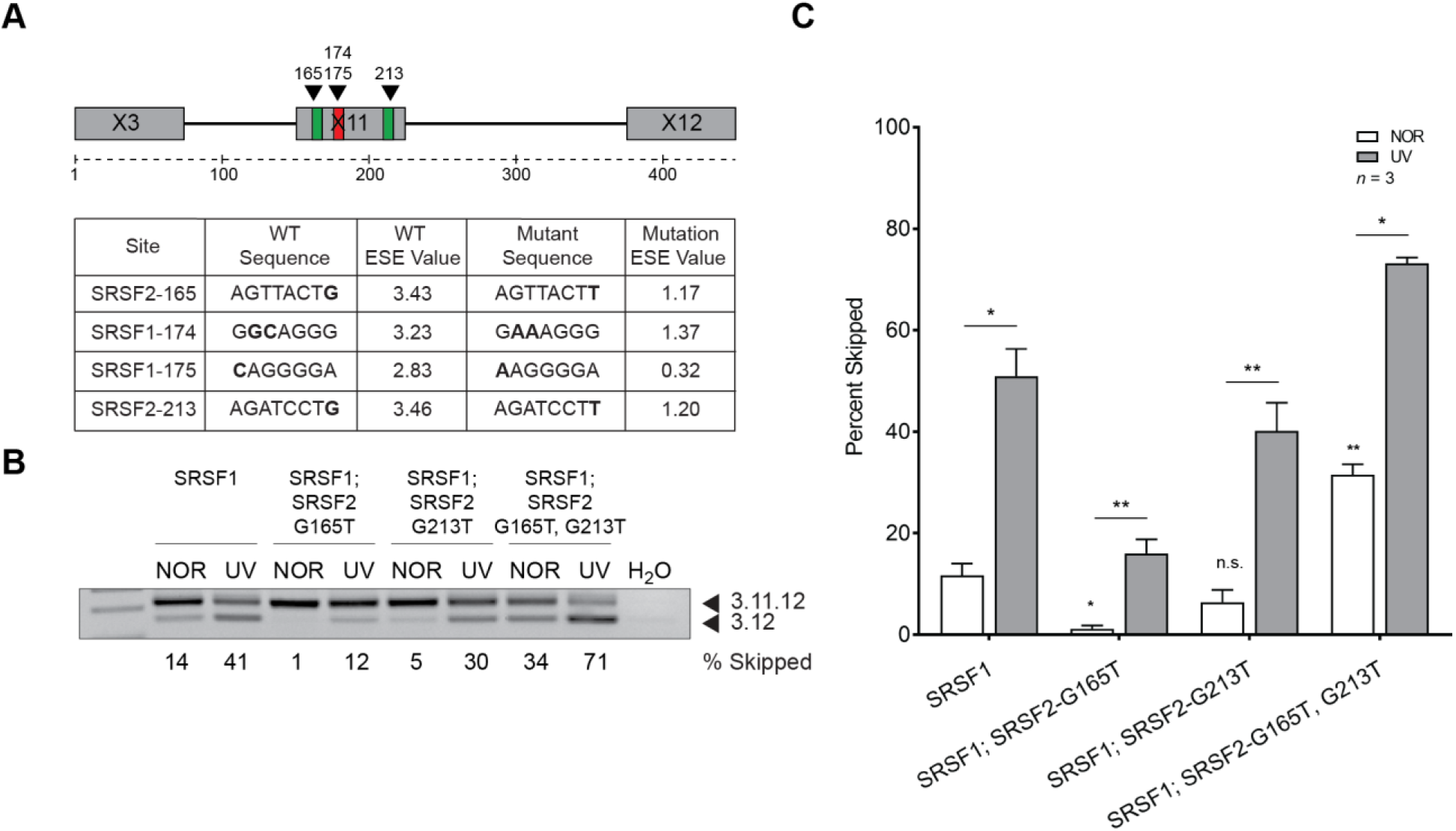
SRSF2 site mutants restore damage-responsive alternative splicing of the *MDM2* 3-11-12s minigene in the presence of SRSF1 site mutants. **A.** Schematic of ESEfinder 3.0-predicted sites for SRSF2 (green), SRSF1 (red), and point mutations made in the *MDM2* 3-11-12s minigene (black triangles). Sequences and matrix scores for wild-type (WT) and mutant exonic splicing enhancer (ESE) values were predicted by ESEfinder 3.0. Mutations made in the sequence of the *MDM2* 3-11-12s minigene lowered the predicted matrix score for binding of splicing factor SRSF2 and SRSF1. **B.** *MDM2* 3-11-12s minigenes were transfected into MCF7 cells for 24 hours, then treated under normal (NOR) or UVC conditions, and harvested 24 hours later. Under normal conditions the three mutant minigenes showed either a reduction (SRSF1; SRSF2-G165, *p* = 0.012), neutral effect (SRSF1; SRSF2-G213T, *p* = 0.192), or increase (SRSF1; SRSF2-G165T, G213T, *p* = 0.003) in expression of the 3.12 skipped product compared to the wild-type minigene. The damage induction of the 3.12 product was maintained in all constructs. **C.** The bar graphs represent the percentage of 3.12 skipped product obtained from three independent experiments under each condition and the error bars represent standard error of the mean (SEM). * indicates *p* < 0.05, ** indicates *p* < 0.01 in all cases as determined by a two-tailed Student’s T test.

## DISCUSSION

We identified SRSF2 as a positive splicing factor that promotes the recognition of exon 11 of MDM2. We demonstrated that the SRSF2 binding sites are conserved between mouse and human *MDM2* exon 11. Furthermore, these sites are sufficient to promote full-length splicing endogenously as splicing is compromised using either the SSOs or CRISPR-Cas9 generated mutations. Exon 11 of *MDM2* is well-conserved between mouse and human. Both are 78 base pairs and share approximately 82% nucleotide identity. While both SRSF2 binding sites are conserved according to *in silico* prediction (20), the first site has a mismatch at 3/8 nucleotides and second is 100% conserved on the sequence level. The absolute requirement for the more potent *MDM2* G213T, *Mdm2* G213A site may explain the pressure for it to remain conserved over evolutionary time. In both mouse and human minigene mutations, as well as in SSO treatment, the second SRSF2 site consistently induced more exon exclusion compared with the first. We also speculate that this is one of the reasons that we were unable to successfully recover a mouse CRISPR cell line with the G213A mutation.

The regulation of SRSF2 is known to be controlled through alternative splicing of its own transcript, as well as posttranslational acetylation and phosphorylation by SRPK1 and SRPK2 (21). Under both normal and UV damage conditions, SRSF2 is localized to nuclear speckles, which are known to be structures where nuclear processing factors are localized, as well as sites of active mRNA transcription (22, 23). In response to UVC treatment we observed that the expression levels of SRSF2 increased and importantly, that the number of speckles to which SRSF2 is localized are fewer in number and have a larger foci diameter. Additionally, we saw that binding of SRSF2 was decreased *in vivo* in response to UV treatment. We infer that the loss of colocalization of SRSF2 with *MDM2* transcripts under conditions of genotoxic stress facilitates *MDM2* alternative splicing.

MDM2 is overexpressed in many types of cancer including osteosarcoma, esophageal cancer, and dedifferentiated liposarcomas (1, 24, 25). These cancers are invariably p53 wild-type and would benefit from persistent alternative splicing of *MDM2* to downregulate MDM2 expression, reactivate p53, and sensitize these tumors to current therapies. Whereas other drugs including nutlin-3a and spiro-oxindole analogs MI-63 and MI-219 have shown potential by disrupting the MDM2-p53 interaction as anticancer strategies (26–28), these particular compounds have not been successful in the clinic likely due to their failure to also inhibit MDMX, another negative regulator of p53. Our lab previously demonstrated that MDM2-ALT1 can bind both full-length MDM2 and MDMX and sequester these proteins in the cytoplasm, thereby acting as a dominant negative to the function of full-length MDM2 or MDMX (3, 29). Promoting the alternative splicing of *MDM2* through the use of SSOs would allow for control of both full-length MDM2 and MDMX activity, thereby elevating the levels of p53 in a cancer cell.

While MDM2 overexpression in cancer is well-documented in the literature, paradoxically *MDM2* alternative splicing has been observed with many types of cancer as well, including bladder (19), colon (30), breast (31), and soft tissue sarcomas such as rhabdomyosarcoma (RMS) (32). We previously reported that *MDM2-ALT1* expression correlated with high-grade disease in RMS and is the most common genetic perturbation in both alveolar and embryonal RMS. Recent data suggests that the mechanisms of MDM2-ALT1-mediated oncogenesis are largely p53 independent as expression of MDM2-ALT1 in a p53-deficient background accelerated tumorigenesis and shifted the observed tumor spectrum *in vivo* (33). Therefore, while it would be beneficial to induce alternative splicing of *MDM2* in cancers where it is overexpressed, in cancers with p53 gain-of-function mutations it may be useful to restore full-length MDM2 to degrade mutant p53. As we have now characterized both positive and negative regulators of *MDM2* alternative splicing, it is plausible that SSO therapy could be used to modulate the levels of p53 through the manipulation of MDM2 alternative splicing. SSOs capable of inducing *MDM2-ALT1* that are potent enough to elicit a biological response should, therefore, be a priority. As current modalities to treat MDM2-overexpressing cancers are limited, the results of our study provide a rationale to develop splicing modulating strategies to modulate the p53 levels in cells and re-sensitize them to chemotherapeutic agents.

## MATERIALS AND METHODS

### Cell culture, growth and transfection conditions

MCF7, NIH 3T3 cells, and C2C12 cells were obtained from ATCC and SMS-CTR cells were obtained from Peter J. Houghton. HeLa S3 cells were obtained from Hua Lou. All human cell lines have been verified by STR analysis (Genetica). Experiments were performed within the first 10 passages of thawing cells. MCF7, NIH 3T3, and C2C12 cell lines were maintained in DMEM, whereas SMS-CTR and HeLa S3 cells were maintained in RMPI medium. Both were supplemented with 10% fetal bovine serum (Catalog Number SH3007103) from Thermo Fisher Scientific, 1X L-glutamine (Catalog Number MT 25-005 CI) from Corning and 1X penicillin/streptomycin (Catalog Number MT 30-001 CI) by Corning. MCF7 cells were transfected with *MDM2* minigenes along with SRSF2 or LacZ overexpression plasmids as previously described (16). For RNA immunoprecipitation assays, MCF7 cells were seeded to 80% confluency in 15 cm plates and transfected with 5.0 μg LacZ or T7-SRSF2 for 24 hours then cells were either treated under normal conditions or subjected to 50 J/m^2^ UVC and harvested 12 hours later.

### Plasmids, protein expression constructs

The LacZ plasmid was previously described (16). The T7-SRSF2 construct was provided as a kind gift from Dr. Adrian Krainer. The *MDM2* 3-11-12s minigene was previous described (16). The SpCas9-2A-EGFP plasmid was obtained from Addgene (Catalog Number 48138). The guide sequences for target sequences were cloned into the BbsI site as previously described (34). The *Mdm2* 3-11-12s minigene was constructed using polymerase chain reaction (PCR) with the following primers: (1F) 5’ GTTCGGATCCGCCAATGTGCAATACCAACATGTCTG 3’ (1R) 5’ TCTCAGTAAGTCTTATGCGATAATCCAGGTTTCAATTTTGTT 3’ (2F) 5’ ACCTGGATTATCGCATAAGACTTACTGAGAATTCTGGCTT 3’ (2R) 5’ GTAACTCGAGCCTCAGCACATGGCTCT 3’. The *Mdm2* 4-11-12s minigene was constructed using polymerase chain reaction (PCR) with the following primers: (1F) 5’ GAGCCCGGGCGGATCCGTTAGACCAAAACCATTGCTTTTGAA 3’ (1R) 5’ TCTCAGTAAGTCTTAATCTCACTCAAACTTGAAAAACCACCA 3’ (2F) 5’ AAGTTTGAGTGAGATTAAGACTTACTGAGAATTCTGGCTT 3’ (2R) 5’ CGGGCCCCCCCTCGAGCCTCAGCACATGGC 3’.PCR products were visualized under longwave ultraviolet, excised, and gel purified using QIAquick Gel Extraction Kit (Catalog Number 28704). A final multiplex PCR was performed with the two purified PCR products and the two terminal primers. The *Mdm2* 3-11-12s minigene was then cloned into the EcoRI-XhoI sites of the pCMV-Tag2B vector using T4 DNA ligase (Catalog No. M0202S) from NEB according to the manufacturer’s instructions.

### RT and PCRs

Reverse transcription (RT) reactions were carried out using 1 μg of RNA using Transcriptor RT enzyme (Catalog No. 03531287001) from Roche Diagnostics. Non-quantitative endogenous *MDM2* PCRs were performed as previously reported (35). *MDM2* minigene PCRs were performed as previously reported (15). PCRs for *MDM2* after RNA immunoprecipitation of T7-SRSF1 were performed using a set of nested primers under the following conditions: (94°C 4’, 35 cycles of 94°C 30′, 62°C 30′, 72°C 1’, 72°C 7’) 5’ TTCCCCTTTACACTCACTT 3’ and 5’ TACAGGTCTCATCACAACAAATAA 3’ then (94°C 5’, 35 cycles of 94°C 30′, 58°C 30′, 72°C 1’, 72°C 7’) 5’ TTTCCCCCTTTACACTCACT 3’ and 5’ AAATTTCAGGATCTTCTTCAA 3’. *Mdm2* amplicons were amplified using the following primers under the standard PCR conditions: (94°C 5’, 35 cycles of 94°C 30′, 55°C 30′, 72°C 1’, 72°C 7’) 5’ TGGCTTCTTGGTTGAAGGGTT 3’ and 5’ CAGCTAAGGAAATCTCAGGATCT 3’.

### Quantification of splicing ratios

Percentages of full-length and exon-excluded products were quantitated using ImageQuant TL (Version 8.1). Significance of the results was assessed using the two-tailed Student’s t-test using GraphPad Prism (Version 6.0).

### Western blot analysis and antibodies

Cell were lysed in NP-40 buffer and equal amounts of protein were loaded in SDS sample buffer onto a sodium dodecyl sulfate-polyacrylamide gel (SDS-PAGE), blotted onto a polyvinylidene difluoride (PVDF) membrane, and analyzed for expression of SRSF2 using either clone 1SC-4F11 (Catalog Number 06-1364), T7-Tag (Catalog Number 69522) from EMD Millipore, or β-Actin clone AC-15 (Catalog Number A5441) from Sigma. Protein sizes were determined using the Precision Plus Protein Dual Color Standards marker (Catalog Number 161-0374) from Life Technologies.

### Microscopy

Cells were seeded on coverslips and either treated under normal conditions or subjected to 50 J/m^2^ UVC. After 12 hours cells were fixed in 4% paraformaldehyde and permeabilized in 0.25% Triton X-100. Cells were then blocked in 10% donkey serum and incubated in an SC35 primary antibody (Catalog Number 556363) in 5% donkey serum at 4°C overnight. Cells were incubated in secondary antibody at room temperature for 1 hour in dark (anti-mIgG Alexa Fluor 488 Thermo Fisher A31571 1:1000). Coverslips were then mounted with 1 drop of Diamond ProLong Antifade with DAPI (Thermo Fisher) and cured overnight at room temperature. Cells were then imaged on a confocal microscope.

### RNA Immunoprecipitation

Cells were scraped from adherent plates in 1 mL of PBS on ice. Suspensions were transferred to Eppendorf tubes and spun down for 30 seconds at 1300 x *g.* PBS was aspirated and cells were lysed in polysome lysis buffer (100 mM KCl, 5mM MgCl_2_, 10mM HEPES, pH 7.0, 0. 5% Nonidet P-40, 100U/mL RNase inhibitor, Halt protease inhibitor) on ice for five minutes. Lysates were centrifuged at 14,000 x *g* for 15 minutes at 4°C and supernatant was transferred to a fresh tube. Approximately 1.5 mg of protein lysate were immunoprecipitated in a 1 mL reaction containing NT2 buffer (50 mM Tris, pH 7.4, 150 mM NaCl, 1 mM MgCl_2_, 0.05% Nonidet p-40), 20 μg of T7-SRSF1 or mIgG isotype control with 15 mM EDTA, pH 8.0 for 10 minutes at room temperature. 100 μl of prewashed Dynal Protein G magnetic beads were then added to the immunoprecipitation reaction for an additional 10 minutes. Immunoprecipitates were washed 3 times with 500 μl NT2 buffer, then twice with 500 μl PBS (containing 100 U/mL RNase inhibitor). IPs were resuspend in 150 μl proteinase K buffer (1.2 mg/mL proteinase K, 1% SDS, 100U/mL RNase inhibitor in NT2 buffer) and incubated 30 minutes at 55°C. Beads were immobilized and supernatant was transferred to fresh Eppendorf tubes, to which 350 μl buffer RLT was added to both IP and 1/100 input samples. A Qiagen RNeasy protocol (Catalog 74106) with DNase digestion was then performed according to manufacturer’s instructions.

### RNA oligonucleotide pull down

RNA probes were synthesized from Integrated DNA Technologies (Coralville, IA, USA) (SRSF2-165 WT ‘UAUCAGGCAGGGGAGAGUGAU’, SRSF2-165 MUT ‘UAUCAGAAAGGGGAGAGUGAU’, SRSF2-213 WT ‘UAUCAGGCAGGGGAGAGUGAU’ SRSF2-213 MUT ‘UAUCAGAAAGGGGAGAGUGAU’). A total of 5 nmol of RNA was modified, conjugated Adipic acid dihydrazide agarose beads (Catalog Number A0802-10ML) from Sigma, and washed as previously described (16). RNA was then incubated in a splicing reaction at 30°C for 40 min, gently mixing every 5 min. Protein-bound beads were washed 3X in Buffer D, then eluted in 40 μl 2X SDS Buffer. Beads were boiled 100°C for 5 min, then spun down 10000 rpm at 4°C for 10 min. Eluates were collected and loaded in equal volume on 10% SDS-PAGE gel, transferred to PVDF membrane, and probed for SRSF2 and β-Actin.

### SRSF2 knockdown

The siRNAs targeting human *SRSF2 (SRSF2* 3’ UTR-siRNA sense, UUGGCAGUAUUGACCUUAUU; *SRSF2* 3’ UTR-siRNA antisense, UAGGUCAAUACUGCCAAUU) or a non-specific siRNA (CTRL sense, AAGGUCCGGCUCCCCCAAAUG; CTRL antisense, CAUUUGGGGGAGCCGGACCUU) were synthesized by Life Technologies. siRNAs were transfected into MCF7 cells at a concentration of 30 nM, mediated by Lipofectamine RNAiMAX from Life Technologies for a total of 72 h. Posttransfection cells were harvested for total RNA using an RNeasy kit (Catalog 74106) from Qiagen and subject to RT-PCR as described above. Protein was also collected as described above to confirm knockdown of SRSF2.

### SSO treatment

2’O-methyl SSOs were provided by Trilink BioTechnologies. SSOs specific to *MDM2* exon 11 (#1 ‘CUGCCUGAUACACAGUAACU’, #2 ‘UUUCAGCAUCUUCUUCAAAU’, #3 ‘GAAAUUUCAGGAUCUUCUUC’) or a non-specific SSO (‘AUAUAGCGACAGCAUCUUCC’) were transfected in MCF7 cells with either Lipofectamine 2000 (Catalog Number 11668030) Lipofectamine 3000 (Catalog Number 15338-100) from Life Technologies for 24 hours. SMS-CTR cells were nucleofected with 250 nM SSO using Nucleofector Kit R (Catolog Number VACA-1001) with program X-001 on an Amaxa Nucleofector II device. All cells were harvested for RNA after 48 hours of transfection using a RNeasy kit from Qiagen and subjected to qPCR using conditions described above.

### Quantitative Real-Time PCR (qPCR)

All Quantitative qPCR was performed with standard PCR conditions for using an Applied Biosystems 7900HT Fast Real Time PCR system (Life Technologies). Real-time PCR reactions were carried out using the SYBR Green PCR master mix (Applied Biosystems part no. 4309155). The primers used to amplify the p53-target transcripts and *MDM2-ALT1* have been previously described (6). *Mdm2-MS2* was amplified using the following primers: 5’ ACACTATGAAAGAGGACTATTGGAA 3’ and 5’ TTTCACGCTTTCTTGGCTGC 3’. All PCR reactions were carried out with 3 technical replicates and the amplification of single PCR products in each reaction was confirmed using dissociation curve.

### CRISPR-Cas9 genome editing

NIH 3T3 were transfected with 1.0 μg of control plasmid SpCas9-2A-EGFP or SpCas9-2A-EGFP-MDM2 along with 500 ng of an HDR custom single-stranded DNA 243 base repair template from IDT DNA. 4 hours after transfection cells were treated with a 1 mM SCR7 (Catalog Number M60082-2s) from Xcessbio. 48 hours after transfection, cells were sorted for GFP expression on a BD Influx FACS cell sorter running SortWare software. GFP-positive cells were plated and maintained in medium containing 1 mM SCR7. After two passages, cells were collected for genomic DNA. Genomic changes were verified by performing PCR of *Mdm2*, followed by TOPO cloning of PCR products, and analyzed by Sanger sequencing (MWG Eurofins).

### Live cellular growth assay

Growth curves were performed with triplicate plating of either NIH 3T3 Control CRISPR, NIH 3T3 G165T CRISPR I or NIH 3T3 G165T CRISPR II cell lines. Cells were seeded at a density of 8 × 10^4^ cells per well in a 12-well plate and analyzed for confluency using the IncuCyte ZOOM™ live cell imaging system taking pictures every 4 hours. Statistical significance was calculated by two-way ANOVA.

## Acknowledgements

We would like to thank Aixa S. Tapia-Santos and Jordan T. Gladman for their technical contribution. We would also like to thank the members of the Chandler lab for critical review of the manuscript.

